# Three genes controlling streptomycin susceptibility in *Agrobacterium fabrum*

**DOI:** 10.1101/2023.05.25.542317

**Authors:** Robyn E. Howarth, Curtis M. Pattillo, Joel S. Griffitts, Diana G. Calvopina-Chavez

## Abstract

Streptomycin is a commonly used antibiotic for its efficacy against diverse bacteria. The plant pathogen *Agrobacterium fabrum* is a model for studying pathogenesis and interkingdom gene transfer. Streptomycin-resistant variants of *A. fabrum* are commonly employed in genetic analyses, yet mechanisms of resistance and susceptibility to streptomycin in this organism have not previously been investigated. We observe that resistance to a high concentration of streptomycin arises at high frequency in *A. fabrum* and we link this trait to the presence of a chromosomal gene (*strB*) encoding a putative aminoglycoside phosphotransferase. We show how *strB*, along with *rpsL* (encoding ribosomal protein S12) and *rsmG* (encoding a 16S rRNA methyltransferase) modulate streptomycin sensitivity in *A. fabrum*.

## IMPORTANCE

The plant pathogen *Agrobacterium fabrum* is a widely used model bacterium for studying biofilms, bacterial motility, pathogenesis, and gene transfer from bacteria to plants. Streptomycin is an aminoglycoside antibiotic known for its broad efficacy against gram-negative bacteria. *A. fabrum* exhibits endogenous resistance to somewhat high levels of streptomycin, but the mechanism underlying this resistance has not been elucidated. Here, we link this resistance to a chromosomally encoded orphan StrB homolog that has not been previously characterized in *A. fabrum*. Furthermore, we show how the genes *rsmG*, *rpsL*, and *strB* jointly modulate streptomycin susceptibility in *A. fabrum*.

## INTRODUCTION

Streptomycin (Sm) inhibits the fidelity of the prokaryotic ribosome by stabilizing a conformational state of the 16S rRNA that results in codon-anticodon mismatches during translation (1). Sm binds the ribosome at an interface between several 16S helices, including helix 18, and the ribosomal protein S12 (**Fig. 1**) (2, 3). Sm resistance has been linked to mutations in *rpsL*, *rsmG* (also known as *gidB*), and *rrs* which, respectively, encode ribosomal protein S12, S-adenosylmethionine (SAM)-dependent 16S rRNA methyltransferase (RsmG), and 16S rRNA.

**FIG 1.**
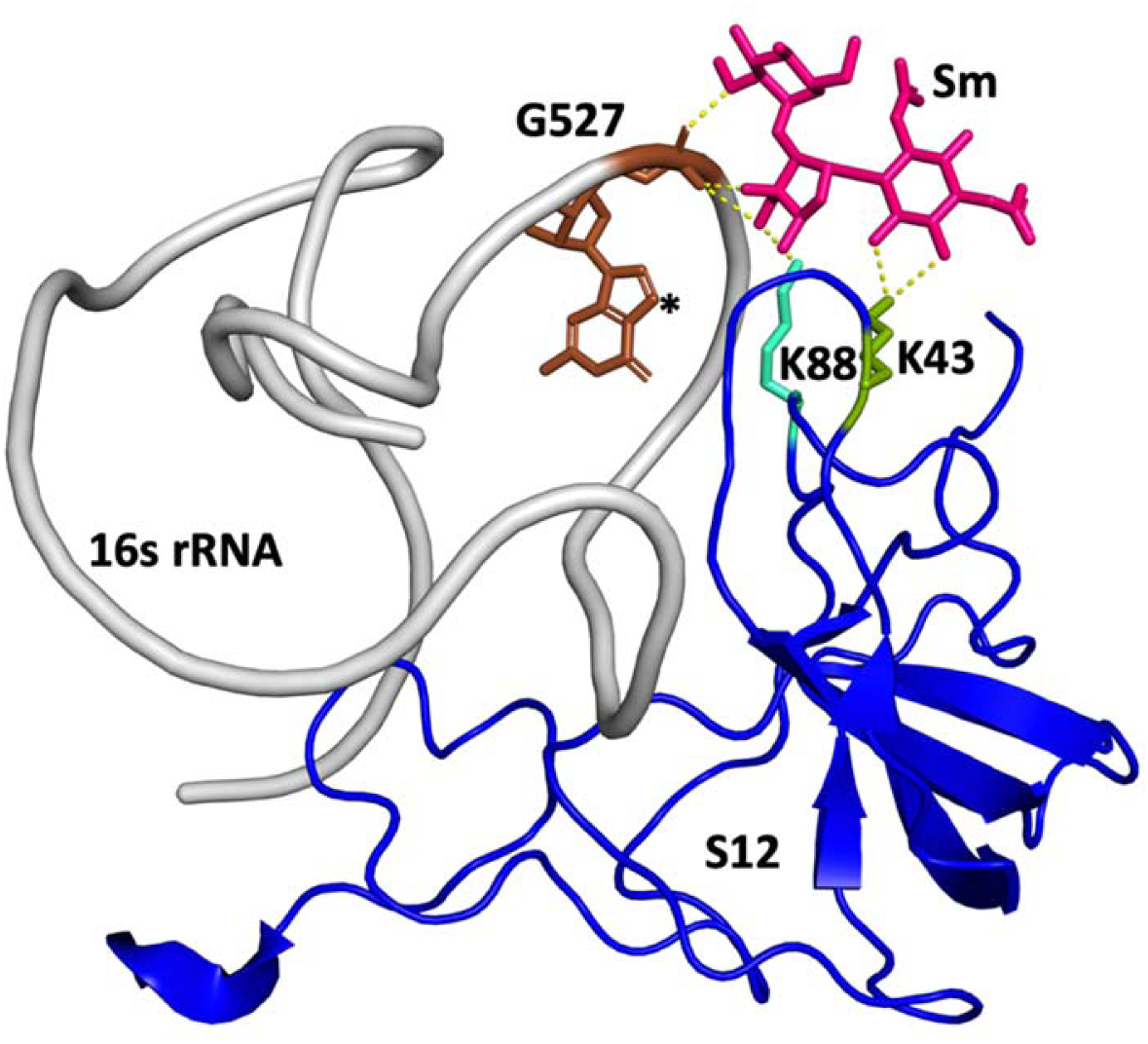
Cartoon representation of a portion of the 30S subunit that shows key interactions with Sm. Sm is shown in pink sticks. Helix 18 of the 16s rRNA is shown in gray with guanosine 527 highlighted in brown. The asterisk shows the nitrogen atom that is methylated by RsmG. Ribosomal protein S12 is shown in blue with key residues K88 and K43 shown in cyan and green respectively.

In diverse bacteria, high-level streptomycin resistance has been linked to point mutations in the S12-encoding *rpsL* gene (4, 5). S12 is located at the interface of the large and small ribosomal subunits, where it interacts with the EF-Tu-bound Trna acceptor arm and functions as a control element for translocation of the mRNA:tRNA complex (6–8). In *Escherichia coli*, spontaneous mutations in the *rpsL* gene that result in a single amino acid change (K42R or K87R) confer high levels of Sm resistance (9, 10). These mutations also occur in Sm-resistant strains of *Mycobacterium tuberculosis* and *Streptomyces coelicolor* (11–13).

RsmG is a member of a large family of SAM-dependent methyltransferases functioning in cell division and chromosome replication. In many bacteria, such as *E. coli* and *Bacillus subtilis*, RsmG has been shown to be responsible for N7 methylation of the 16S rRNA at position G527 located on the highly conserved helix 18 (**Fig. 1**) (14, 15). Sm has been shown to interact with the phosphate backbone of G527 (3, 16). In *S. coelicolor, M. tuberculosis*, *B. subtilis*, and *Thermus thermophilus*, loss of *rsmG* results in low-level Sm resistance likely due to the loss of this key methylation event occurring near the Sm binding pocket (14, 15, 17–19).

Alterations in the 16S sequence are generally not associated with Sm resistance because most bacteria possess many redundant copies of the 16S-encoding gene (*rrs*), making any single *rrs* mutation recessive. However, mutations in the *rrs* gene that confer Sm resistance can be found by genetically modifying bacteria to carry a single functional copy of *rrs* and selecting for Sm-resistant mutants. For example, in *M. smegmatis*, mutations in the *rrs* gene were selected by altering the number of *rrs* alleles in the bacterial genome. Most of the mutations mapped to the highly conserved 530 loop region of the 16S rRNA, specifically the mutation 524G>C which has been thought to be essential for ribosome function (20).

Sm resistance may also be brought about by inactivating enzymes. The *strA-strB* resistance cassette has been characterized in taxonomically diverse Gram-negative bacteria. StrA is an aminoglycoside-3’’-phosphotransferase and StrB is an aminoglycoside-6-phosphotransferase (21). The pair of enzymes is thought to work in concert to inactivate Sm, with loss of either gene being associated with loss of strong resistance (22–25).

The plant pathogen, *Agrobacterium fabrum* C58, has become an important model for studying interkingdom gene transfer, cell polarity, and motility (26, 27). Here, using a plasmid-free derivative strain, we report that *A. fabrum* C58 has moderate Sm resistance due to an unusual chromosomal copy of *strB* without an accompanying *strA* companion gene. In this context, we show how Sm susceptibility is controlled in *A. fabrum* by the *strB*, *rsmG*, and *rpsL* gene network.

## RESULTS

### Frequency and mechanism of Sm resistance in *A. fabrum* varies by Sm concentration

A plasmid-free derivative of *A. fabrum* C58 (UBAPF2) (28) was found to give rise to surprisingly large numbers of Sm-resistant colonies when selected at 200 µg/ml Sm, with an average frequency of 7.1×10^-5^± 2.3×10^-5^ (SD; n=10). However, at 800 μg/ml, colonies emerged over 100 times less frequently, with an average frequency of 4.3×10^-7^ ± 2.1×10^-7^ (SD; n=10). Each culture in these analyses was derived from an independent colony in order to account for fluctuation in the data. We sequenced *rpsL* for several Sm^200^- and Sm^800^-resistant derivatives and found sequence changes only in the Sm^800^ group (with a major allele being K43R), suggesting that the mechanism of resistance for Sm^200^ derivatives is not mediated by *rpsL*.

To determine the genetic basis of resistance in Sm^200^ derivatives, whole-genome resequencing was carried out on six independent isolates. In each of the isolates, a mutation was found in *rsmG* (*ATU2830*, also known as *gidB*), and these are depicted on the map in **Fig. 2**. These *rsmG* alleles are mostly predicted to be associated with loss of function due to frameshift or nonsense mutations. Aside from these six mutations in *rsmG*, only two additional sequence deviations from the reference genome were identified across the six resequenced strains: one intergenic substitution in strain YS01, and one missense mutation (Gly to Ala) in the riboflavin biosynthesis *ribB* gene in strain BB01. From this, we conclude that Sm^200^ resistance in these UBAPF2 derivatives was brought about by the observed changes in *rsmG*.

**FIG 2.**
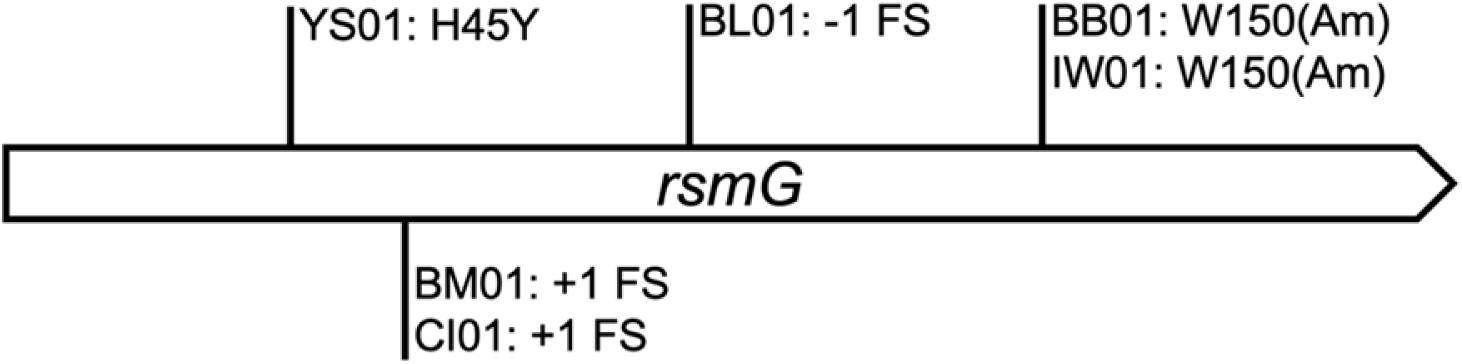
Spontaneous mutant alleles of *rsmG* associated with Sm^200^ resistance in *A. fabrum*. Six independently derived isolates (YS01, BM01, CI01, BL01, BB01, and IW01) selected on Sm were whole-genome sequenced and each exhibited a mutation in *rsmG* (FS, frameshift; Am, premature amber stop codon).

### *strB* provides background Sm resistance in *A. fabrum*

For many Gram-negative, Sm-sensitive bacteria, Sm^200^ is considered a high dose. In *E. coli* K12 for example, we observe resistance to Sm^200^ to occur at a frequency of less than 1×10^-9^, and the mechanism is uniformly *rpsL*-mediated (data not shown). Mutations in *rsmG* are generally associated with low-level Sm resistance (14, 15, 17). This suggests that the parental UBAPF2 strain possesses significant background resistance. Investigating this further, we found the minimal inhibitory concentration (MIC) of Sm be around 128 µg/ml for UBAPF2. We hypothesized that this native-level resistance is caused by an endogenous, dominantly acting gene. We reasoned that random chromosomal insertion of a strong promoter could help us to identify this factor by screening for elevated Sm resistance. The Tn5-110 transposon carries the outwardly oriented P_trp_ promoter from *Salmonella* (29) that has successfully given overexpression phenotypes in *Sinorhizobium meliloti*. We conjugated the Tn5-110 delivery plasmid into UBAPF2 and selected for growth on Sm^200^ plates (additionally containing neomycin to select for transposon insertion). Fifteen colonies from this selection were evaluated for transposon insertion location. In 7 of these, insertions were distributed around the genome with no clear pattern (Table S1); in the other 8, the insertions occurred in varying positions within a small genomic interval, shown in **Fig. 3**. These 8 insertions all position the P_trp_ promoter in the same orientation, reading into a pair of likely co-transcribed genes: ATU1244, and ATU1243. ATU1244 (*argC*) encodes a N-acetyl-gamma-glutamyl-phosphate reductase enzyme predicted to be involved in the biosynthesis of arginine and ornithine. Downstream, ATU1243 (*strB*) encodes an StrB family phosphotransferase, possibly involved in modification of streptomycin or similar aminoglycoside antibiotics. This gene has not been previously associated with Sm resistance in *A. fabrum*. Considering that *strB* genes are usually linked to *strA* partner genes (21, 23, 30), we sought to identify potential *strA* homologues in *A. fabrum* C58. In a BlastP search using the canonical StrA/StrB protein sequences encoded by *E. coli* plasmid RSF1010 (22, 24), we identified the *A. fabrum strB* gene reported above, but no reported homologue for *strA*. The *A. fabrum strB* gene resides in a genomic region that is generally conserved across many species in the Rhizobiaceae family. For example, the *speB-argC* gene pair found upstream of *strB* is well conserved in this family, as well as nearby ribosomal protein genes *rpsI and rplM*; however interspecies comparison of this genomic region shows that it is punctuated by certain species-specific genes (**Fig. 4**). This comparison indicates that *strB* is one of the non-conserved genes in this region, found in some *Agrobacterium* species but absent in most of the Rhizobiaceae genera that we have evaluated.

**FIG 3.**
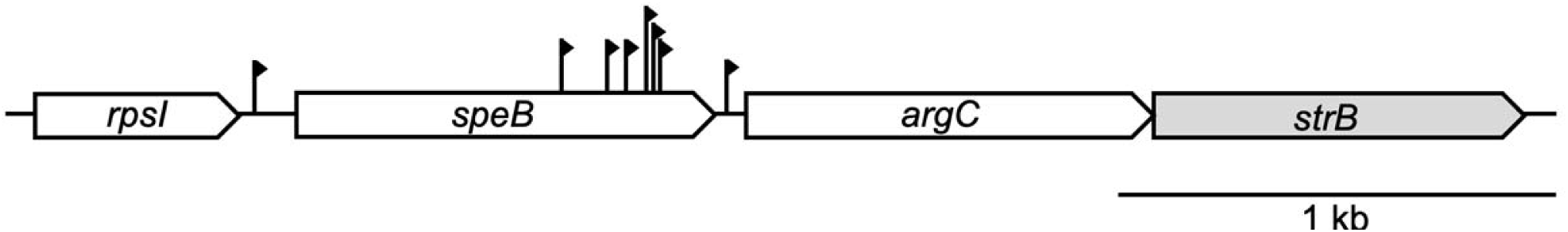
Tn5-110 transposon insertions giving rise to Sm^200^ resistance in *A. fabrum*. Mapped insertion sites are indicated by vertical lines. Direction of transcription from the strong P_trp_ promoter on the transposon is indicated by filled arrowheads. The *strB* gene suspected of being required for this resistance is highlighted in gray.

**FIG 4.**
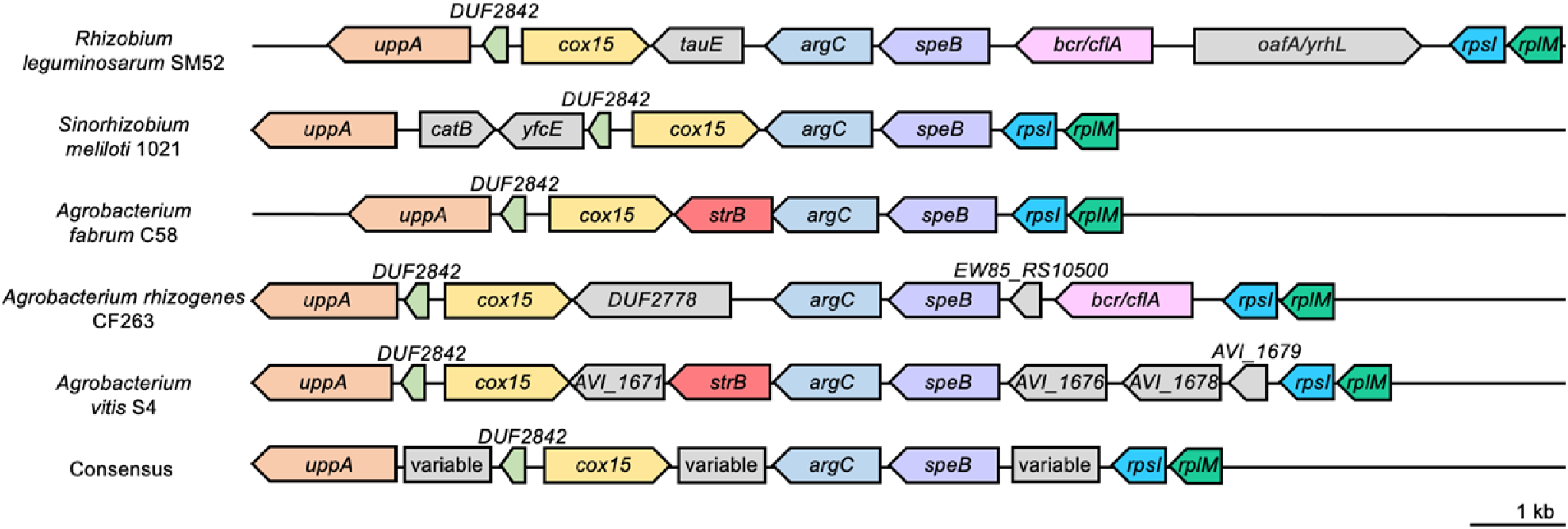
Interspecies comparison of the *strB* genomic region. Conserved genes across species are depicted in the same colors while variable genes are shown in gray. The *strB* gene is depicted in red.

We reasoned that deletion of *strB* would significantly reduce the Sm resistance observed for our parental strain. To test this, *strB* was removed using allele exchange, leaving only the first and last ten codons of the gene intact. This deletion strain was found to have over 60-fold greater sensitivity to Sm, with an MIC of 2 μg/ml (see **Fig. 5** below). In this genetic background, we found that Sm-resistant colonies arise at low frequency (approximately 5×10^-7^) on both 200 µg/ml and 800 µg/ml Sm, suggesting that *rpsL*-mediated resistance is the predominant mechanism. Indeed, all colonies analyzed from these selections (4/4 for 200 µg/ml and 4/4 for 800 µg/ml) harbored *rpsL* mutations. Six of these had the K43R allele, and two of the Sm^200^-resistant mutants had the K88R allele.

**FIG 5.**
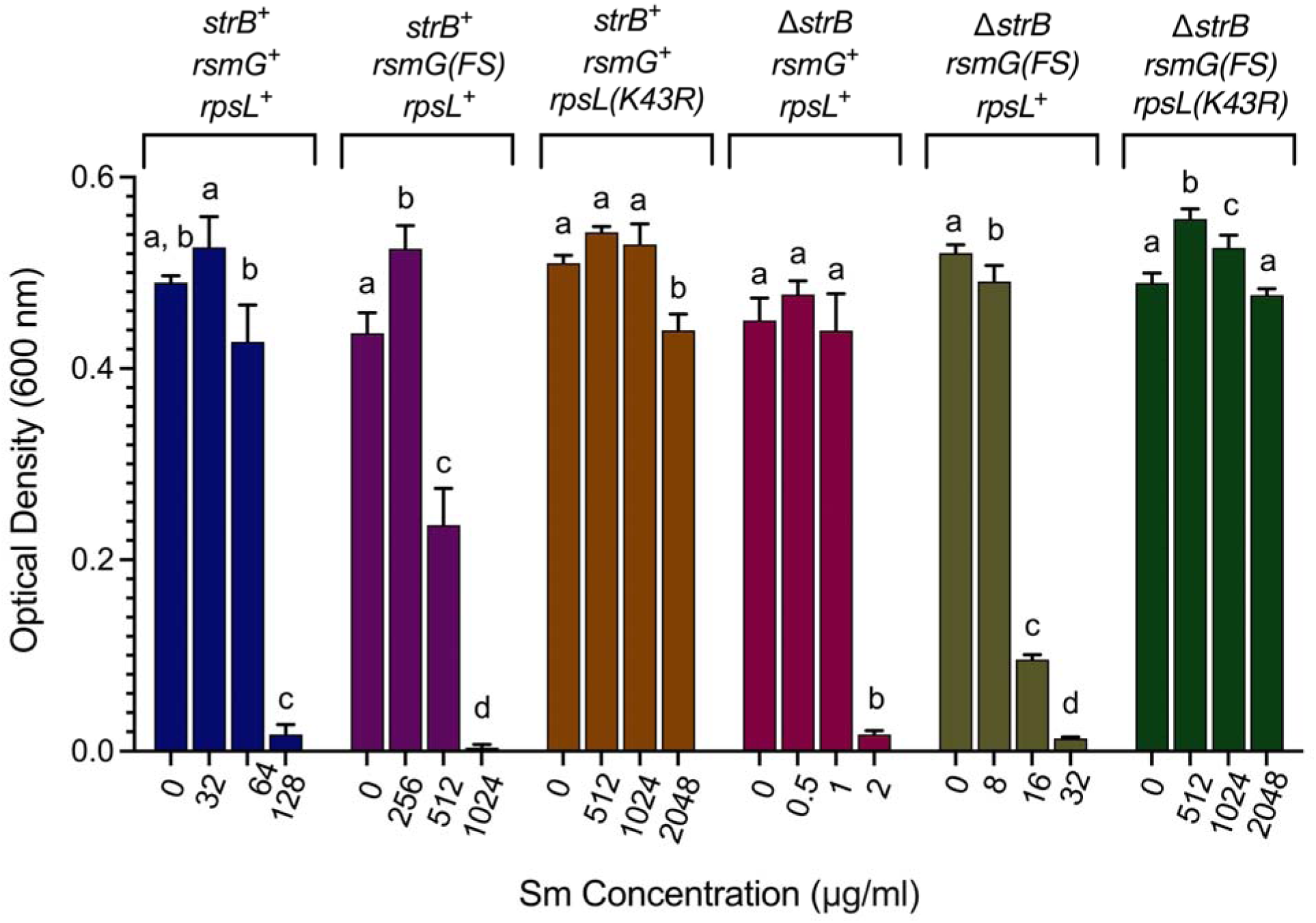
Sm dose responses for six *A. fabrum* genotypes tested. Optical density measurements were taken 20 hours after inoculation of 200-µl cultures in 96-well plates. Genotype descriptions are given above each set of growth values. Error bars represent standard deviation from the mean (n=3). Different letters denote statistically significant differences (P<0.05) according to a Tukey multiple comparison test.

To test the sufficiency of *strB* to confer Sm resistance, it was ligated into a small constitutive expression plasmid and tested for its ability to provide Sm resistance to *E. coli*. As shown in **Fig. 6**, this plasmid allowed growth of *E. coli* up to 160 µg/ml Sm, whereas the vector-only control strain was unable to grow at all non-zero doses tested. The strain expressing *strB* exhibited a significant growth defect in the absence of Sm, likely a result of a metabolic cost from constitutive expression of this resistance gene.

**FIG 6.**
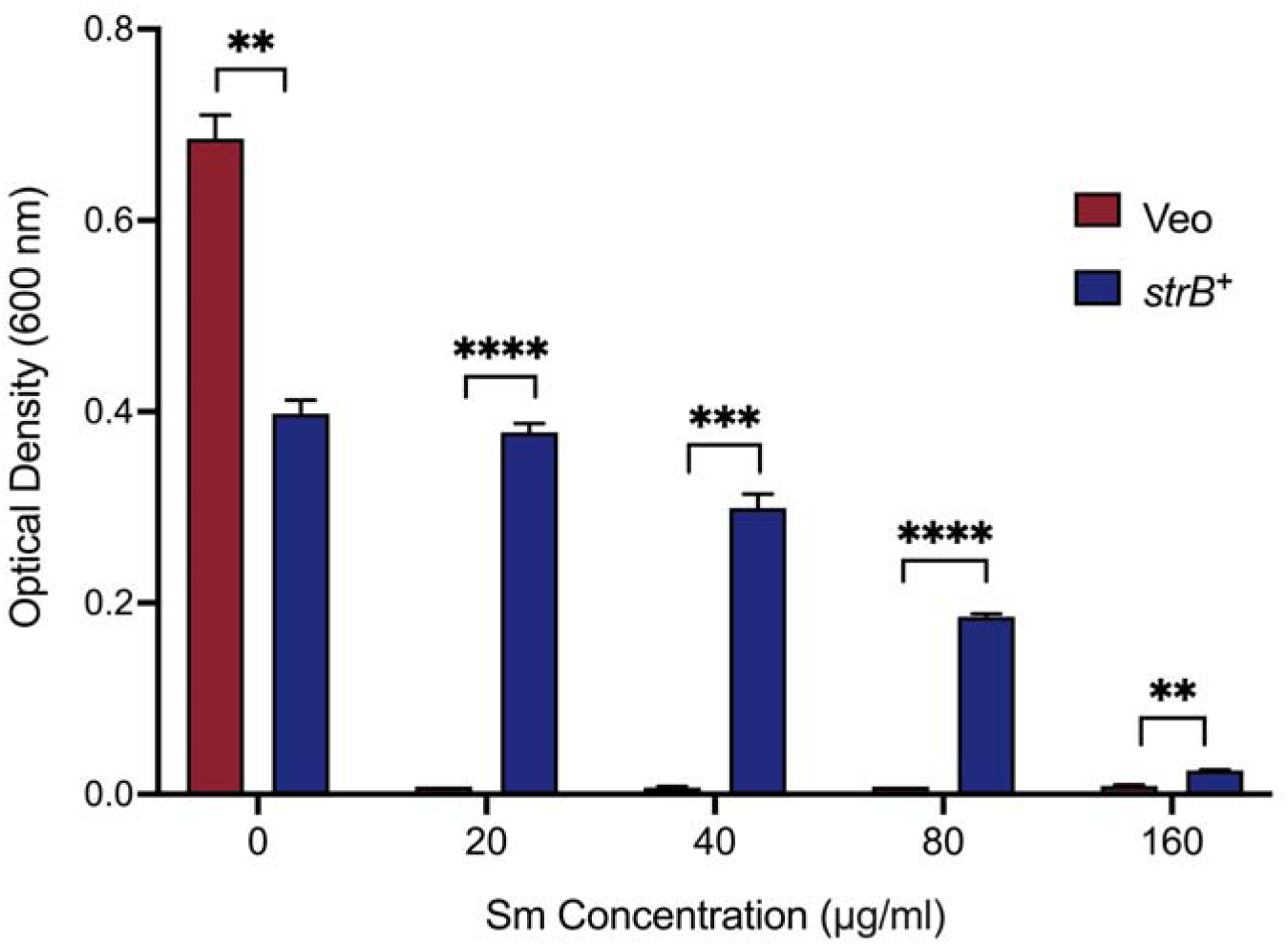
Dose responses for two strains of *E. coli* in the presence of Sm. The *strB*^+^ strain (blue) contains a plasmid that constitutively expresses *A. fabrum strB* while the Veo strain (red) contains the empty parent vector. Error bars show standard deviation from the mean (n=3). Significant differences (\*\*\*\**P* < 0.000001; \*\*\**P* < 0.000005; \*\**P* < 0.00007) are indicated by asterisks according to parametric t-tests carried out with the Benjamín, Krieger and Yekutieli method.

### *rpsL*, *rsmG*, and *strB* constitute a three-gene network modulating Sm resistance in *A. fabrum*

The results outlined thus far point to a model in which three different *A. fabrum* genes influence Sm sensitivity: *strB* provides a modest level of resistance by inactivation of the antibiotic, *rsmG* loss of function can boost resistance by subtly altering the Sm binding site on the ribosome without greatly affecting strain fitness, and very rare and specific mutations in the essential *rpsL* gene can confer greatly elevated resistance due to binding site alteration. This model predicts that the impact of *rsmG* loss of function is strongly modulated by the presence or absence of *strB*, but that Sm resistance-associated *rpsL* mutations provide very strong resistance whether *strB* is present or not. Six genotypes were constructed to test this: i) *strB^+^ rsmG^+^ rpsL^+^*, ii) *strB^+^ rsmG(FS) rpsL^+^,* iii) *strB^+^ rsmG^+^ rpsL(K43R)*, iv) Δ*strB rsmG^+^ rpsL^+^*, v) Δ*strB rsmG(FS) rpsL^+^,* and vi) Δ*strB rsmG^+^ rpsL(K43R)*. Sm dose-response data for these six strains are given in **Fig. 5**. We see from this that tolerance to Sm across all six strains was consistent with our model. The *rpsL(K43R)* allele confers extremely high resistance, whether or not *strB* is intact; and *rsmG* loss of function modestly enhances resistance in the presence or absence of *strB*. Remarkably, *rsmG* loss of function increases resistance by a similar factor (∼10-fold) in the presence or absence of *strB*, indicating that the influence of each gene on resistance is independent and additive.

## DISCUSSION

In this study, three genes were found to have an effect on *A. fabrum* resistance to Sm. A chromosomal *strB* homolog provides moderate resistance which can be enhanced by mutations in either *rsmG* or *rpsL*, the former yielding resistance at higher frequency and the latter allowing resistance at higher doses. The marked difference in frequency of resistance brought about by changes in *rsmG* compared to *rpsL* may be explained by the essentiality of *rpsL* function for cell viability, and so only special alleles of *rpsL* can support both viability and resistance (9). The *rsmG* gene, on the other hand, does not appear to be essential for viability, though deficiency in this gene is associated with only modest resistance to Sm (14, 15, 17). K43R and K88R missense mutations in *rpsL* have been correlated with high-level Sm resistance in several species of bacteria including *Yersinia pestis* (31) and *M. tuberculosis* (32), indicating the conserved molecular-level conservation of Sm binding to this region of the ribosome, and the narrow spectrum of allelic variants of *rpsL* that can support both viability and Sm resistance.

Two-gene *strA-strB* cassettes are commonly associated with Sm inactivation via phosphorylation. Orphan *strA*-only or *strB*-only loci are rarely encountered in bacteria and have not been demonstrably associated with Sm resistance. In *Erwinia amylovora*, Sm resistance was found to significantly decrease with the deletion of either component of its plasmid-encoded *strA-strB* cassette (21). Our observations relating to *strB* in *A. fabrum* are notable in two respects: first is that it is not associated with an *strA* homolog, and second, that it is located on the chromosome rather than on a plasmid. Across five closely-related species, *(Rhizobium leguminosarum*, *S. meliloti*, *A. fabrum*, *A. rhizogenes*, and *A. vitis*) an alignment analysis was performed on the region around *strB*. In all of these species, the genes *uppA*, *DUF2842*, *cox15*, *argC*, *speB*, *rpsI*, and *rplM* are conserved **(Fig. 4)**, but *strB* appears only in the two *Agrobacterium* species (*A. fabrum* and *A. vitis*). The *strB* location (between *cox15* and *argC*) appears to harbor variable genes in the Rhizobiaceae family members we analyzed.

## MATERIALS AND METHODS

### Bacterial strains and growth conditions

*A. fabrum* and *E. coli* strains were grown in Luria Broth (LB) containing (per liter) 10 g tryptone, 5 g yeast extract, 5 g NaCl and 1 ml of 2N NaOH, with 12 g of agar added to solidify when appropriate. *A. fabrum* was grown at 30°C for 2 days while *E. coli* was grown overnight at 37°C. Where appropriate, antibiotics were used as follows: streptomycin (Sm), typically 200 μg/ml or 800 µg/ml; chloramphenicol (Cm), 30 μg/ml; kanamycin (Km), 30 μg/ml; and neomycin (Nm), 100 μg/ml; and rifampicin (Rf), 100 μg/ml. When needed, LB was supplemented with 100 μg/ml of 5-Bromo-4-chloro-3-indoxyl-beta-D-glucuronide cyclohexylammonium salt (X-Gluc) and 1% sucrose. All strains and plasmids used in this study are given in Tables S2 and S3, respectively. Primer sequences are given in Table S4. Relevant plasmid sequences are given in Supplemental Materials.

### Calculating the frequency of spontaneous mutations

To evaluate the tendency of our starting strain UBAPF2 to mutate to Sm resistance, ten independent colonies were grown to saturation in separate liquid cultures. From these, cells were plated on Sm^800^ (800 µg/ml), Sm^200^ (200 µg/ml) or no-Sm LB plates. From colony counts, mean frequencies (Sm^R^/Total) and standard deviation values were calculated.

### Selection of streptomycin-resistant mutants and *rpsL* Sanger sequencing

Six independent Sm^200^-resistant *A. fabrum* colonies were stablished as strains BB01, BL01, BM01, CI01, IW01, and YS01. The *rpsL* gene was amplified from each. PCR was carried out under standard conditions using Taq polymerase and primers 2235 and 2236. Lysed cells, used as template for PCR, were prepared by suspending cells in 200 μl of PCR lysis buffer (5 mM Tris pH 8.0, 2 mM EDTA, 0.5% Triton X-100) and heating to 95°C for 5 min with intermittent vortexing. PCR products were purified using the ZR Plasmid Miniprep-Classic Kit (Zymo Research), followed by Sanger sequencing using either primer 2235 or 2236.

### Whole-genome re-sequencing

Whole-genome sequencing was performed for the six Sm-resistant strains listed above, as well as the wild-type parent strain. Genomic DNA samples were produced with a final concentration greater than 20 ng/µl, using Proteinase K-mediated lysis followed by column purification using the DNeasy PowerLyzer Microbial Kit (Qiagen). These seven samples were sent to Microbial Genome Sequencing Center (MiGS) for Illumina sequencing to return 200 Mbp of data per strain. Resulting FASTQ files were inputted with the GenBank file for the C58 strain of *A. fabrum* into the command-line tool, *breseq*, using Windows Subsystem for Linux and R (33). The *breseq* output allowed for discernment of sequence variants compared to the reference.

### Transposon mutagenesis and determination of insertion sites

A large-scale transposon mutagenesis was performed on *A. fabrum* using pJG110, the delivery plasmid for transposon Tn5-110 (29). A triparental mating was carried out to mobilize the transposon delivery plasmid into *A. fabrum*. This was done by combining the wild-type *A. fabrum* strain (UBAPF2; recipient), the donor strain (DH5α-pJG110) and the helper strain (B001) into a mixed suspension, plating mixed cells onto plain LB, and incubating at 30°C for 24 hours. Resulting lawns were resuspended, plated onto LB-agar containing Sm (200 µg/ml) and Nm, and incubated at 30°C to select for transposants with Sm resistance. 24 medium and large colonies were analyzed by arbitrary-PCR to determine transposon insertion sites. Bacterial template DNA for arbitrary-PCR was prepared by cell lysis as described above. Arbitrary-PCR was carried out as described by Calvopina-Chavez et al. (34), except primers 2133 and 2135 were used for the first-round PCR, and primers 2134 and 2137 were used for the second-round PCR. DNA products were purified as described above and Sanger sequenced using primer 2134.

### Construction of the **Δ***strB* strain D272

Allelic exchange plasmid pJG1108 (34), containing the *gus* and *sacB* genes, was used for *strB* deletion in UBAPF2. Primers oDC103 and oDC104 were designed to amplify the *strB* left homology region, and oDC105 and oDC106 were deigned to amplify the right homology region. The two fragments were amplified by PCR using the High-fidelity Q5 polymerase and they were inserted into XbaI/SalI-digested pJG1108 in a three-fragment ligation. Cloned inserts were then amplified and sequence-verified using primers CD49 and CD50. The resulting *strB* knock-out plasmid (pJG1197) was conjugated into *A. fabrum* UBAPF2 via triparental mating as described above for transposon mutagenesis, though in this case, single cross-over transconjugants were selected on Rf and Nm. Subsequent selection for plasmid eviction was carried out on LB containing X-Gluc and Sucrose. For several resultant white colonies, deletion of *strB* was evaluated by colony PCR using Taq polymerase and primers oDC107 and oDC108. Products were then Sanger sequenced to confirm the deletion.

### Creating a plasmid for constitutive expression of *strB*

Parent plasmid pJG1226 consists of a p15A origin and a Cm resistance gene expressed from a constitutive P_trc_ promoter (pJG1226; sequence is given in Supplemental Materials). A segment with these elements was amplified from pJG1226 with primers oDC196 and oDC197 and digested with XbaI and HindIII. The *strB* gene was amplified from UBAPF2 genomic DNA with primers oDC198 and oDC199, and also digested with XbaI and HindIII. Ligation of the two fragments (pDC76) places *strB* immediately downstream of the *cat* (Cm^R^) gene such that the two are co-transcribed.

### Testing *strB*-dependent Sm resistance in *E. coli*

*E. coli* strain DH5α harboring either pDC76 or pJG1226 (vector-only) was grown in 5 ml of LB+Cm at 37°C overnight. 5 µl of overnight culture was added to 5 ml of LB+Cm containing 0, 20, 40, 80 or 160 µg/ml Sm in triplicates and allowed to grow for 6 h at 37°C. Optical density of each culture was measured at a wavelength of 600 nm, after which data were plotted. Unpaired parametric t-tests were carried out with the Benjamín, Krieger and Yekutieli method to determine statistically significant differences.

### Construction of allelic combinations of *strB*, *rsmG* and *rpsL*

To learn how *strB*, *rsmG*, and *rpsL* interact to modulate Sm resistance in *A. fabrum*, six strains with different allelic combinations of these three genes were assessed: UBAPF2 (*strB^+^ rsmG^+^ rpsL^+^*), D337 (*strB^+^ rsmG(FS) rpsL^+^*), D338 (*strB^+^ rsmG^+^ rpsL(K43R*)), D272 (Δ*strB rsmG^+^ rpsL^+^*), D339 (Δ*strB rsmG(FS) rpsL^+^*), D340 (Δ*strB rsmG^+^ rpsL(K43R*)). The *rsmG(FS)* and *rpsL(K43R)* alleles arise spontaneously with sufficient frequency that they could be introduced by selection on Sm followed by sequence verification. The *rsmG(FS)* allele is a +1 frameshift identical to the allele found in strains BM01 and CI01 (in a homopolymeric run around nt 177 of the *rsmG* coding sequence).

### Determining minimal inhibitory concentrations (MICs)

MICs were determined by growing each strain in triplicate in a 96-well plate containing LB+Sm at appropriate concentrations. The six strains described in the previous paragraph were tested in three concentrations of Sm and a no-Sm control. Specific concentrations are given in **Fig. 5**. For these tests, each well contained 190 μl of LB, and was inoculated with 10 μl of a 10^-1^ dilution of saturated overnight culture. For each strain tested, Sm concentrations were chosen so that the MIC could be discerned. 96-well plates were shaken for 20 hours at 30°C, and culture densities were assessed by OD measurement at 600 nm. For each strain grown under the four conditions, a one-way ANOVA was carried out using Tukey’s multiple comparisons test.

## DATA AVAILABILITY

All relevant data are within the paper and its Supplemental Material file.

## ACKNOWLEDGEMENTS

This project was financially supported by The National Science Foundation (grant IOS-1755454 to J.S.G.).

## REFERENCES

1. Abad JP, Amils R. 1994. Location of the streptomycin ribosomal binding site explains its pleiotropic effects on protein biosynthesis. J Mol Biol 235:1251–60.

2. Carter AP, Clemons WM, Brodersen DE, Morgan-Warren RJ, Wimberly BT, Ramakrishnan V. 2000. Functional insights from the structure of the 30S ribosomal subunit and its interactions with antibiotics. Nature 407:340–8.

3. Demirci H, Murphy Ft, Murphy E, Gregory ST, Dahlberg AE, Jogl G. 2013. A structural basis for streptomycin-induced misreading of the genetic code. Nat Commun 4:1355.

4. Finken M, Kirschner P, Meier A, Wrede A, Bottger EC. 1993. Molecular basis of streptomycin resistance in *Mycobacterium tuberculosis*: alterations of the ribosomal protein S12 gene and point mutations within a functional 16S ribosomal RNA pseudoknot. Mol Microbiol 9:1239–46.

5. Funatsu G, Nierhaus K, Wittmann HG. 1972. Ribosomal proteins. XXXVII. Determination of allelle types and amino acid exchanges in protein S12 of three streptomycin-resistant mutants of Escherichia coli. Biochim Biophys Acta 287:282–91.

6. Stark H, Rodnina MV, Wieden HJ, Zemlin F, Wintermeyer W, van Heel M. 2002. Ribosome interactions of aminoacyl-tRNA and elongation factor Tu in the codon-recognition complex. Nat Struct Biol 9:849–54.

7. Yusupov MM, Yusupova GZ, Baucom A, Lieberman K, Earnest TN, Cate JH, Noller HF. 2001. Crystal structure of the ribosome at 5.5 A resolution. Science 292:883–96.

8. Demirci H, Wang L, Murphy FVt, Murphy EL, Carr JF, Blanchard SC, Jogl G, Dahlberg AE, Gregory ST. 2013. The central role of protein S12 in organizing the structure of the decoding site of the ribosome. RNA 19:1791–801.

9. Timms AR, Steingrimsdottir H, Lehmann AR, Bridges BA. 1992. Mutant sequences in the *rpsL* gene of *Escherichia coli* B/r: mechanistic implications for spontaneous and ultraviolet light mutagenesis. Mol Gen Genet 232:89–96.

10. Hosaka T, Tamehiro N, Chumpolkulwong N, Hori-Takemoto C, Shirouzu M, Yokoyama S, Ochi K. 2004. The novel mutation K87E in ribosomal protein S12 enhances protein synthesis activity during the late growth phase in *Escherichia coli*. Mol Genet Genomics 271:317–24.

11. Sreevatsan S, Pan X, Stockbauer KE, Williams DL, Kreiswirth BN, Musser JM. 1996. Characterization of *rpsL* and *rrs* mutations in streptomycin-resistant *Mycobacterium tuberculosis* isolates from diverse geographic localities. Antimicrob Agents Chemother 40:1024–6.

12. Okamoto-Hosoya Y, Hosaka T, Ochi K. 2003. An aberrant protein synthesis activity is linked with antibiotic overproduction in *rpsL* mutants of *Streptomyces coelicolor* A3(2). Microbiology (Reading) 149:3299–3309.

13. Khosravi AD, Etemad N, Hashemzadeh M, Khandan Dezfuli S, Goodarzi H. 2017. Frequency of rrs and rpsL mutations in streptomycin-resistant *Mycobacterium tuberculosis* isolates from Iranian patients. J Glob Antimicrob Resist 9:51–56.

14. Nishimura K, Johansen SK, Inaoka T, Hosaka T, Tokuyama S, Tahara Y, Okamoto S, Kawamura F, Douthwaite S, Ochi K. 2007. Identification of the RsmG methyltransferase target as 16S rRNA nucleotide G527 and characterization of *Bacillus subtilis rsmG* mutants. J Bacteriol 189:6068–73.

15. Okamoto S, Tamaru A, Nakajima C, Nishimura K, Tanaka Y, Tokuyama S, Suzuki Y, Ochi K. 2007. Loss of a conserved 7-methylguanosine modification in 16S rRNA confers low-level streptomycin resistance in bacteria. Mol Microbiol 63:1096–106.

16. Powers T, Noller HF. 1991. A functional pseudoknot in 16S ribosomal RNA. EMBO J 10:2203–14.

17. Nishimura K, Hosaka T, Tokuyama S, Okamoto S, Ochi K. 2007. Mutations in *rsmG*, encoding a 16S rRNA methyltransferase, result in low-level streptomycin resistance and antibiotic overproduction in *Streptomyces coelicolor* A3(2). J Bacteriol 189:3876–83.

18. Wong SY, Lee JS, Kwak HK, Via LE, Boshoff HI, Barry CE, 3rd. 2011. Mutations in gidB confer low-level streptomycin resistance in Mycobacterium tuberculosis. Antimicrob Agents Chemother 55:2515–22.

19. Gregory ST, Demirci H, Belardinelli R, Monshupanee T, Gualerzi C, Dahlberg AE, Jogl G. 2009. Structural and functional studies of the *Thermus thermophilus* 16S rRNA methyltransferase RsmG. RNA 15:1693–704.

20. Springer B, Kidan YG, Prammananan T, Ellrott K, Bottger EC, Sander P. 2001. Mechanisms of streptomycin resistance: selection of mutations in the 16S rRNA gene conferring resistance. Antimicrob Agents Chemother 45:2877–84.

21. Chiou CS, Jones AL. 1995. Expression and identification of the strA-strB gene pair from streptomycin-resistant *Erwinia amylovora*. Gene 152:47–51.

22. Petrova MA, Gorlenko Zh M, Soina VS, Mindlin SZ. 2008. Association of the strA-strB genes with plasmids and transposons in the present-day bacteria and in bacterial strains from permafrost. Genetika 44:1281–6.

23. Sundin GW, Bender CL. 1996. Dissemination of the strA-strB streptomycin-resistance genes among commensal and pathogenic bacteria from humans, animals, and plants. Mol Ecol 5:133–43.

24. Sundin GW. 2002. Distinct recent lineages of the strA-strB streptomycin-resistance genes in clinical and environmental bacteria. Curr Microbiol 45:63–9.

25. Kim C, Mobashery S. 2005. Phosphoryl transfer by aminoglycoside 3’-phosphotransferases and manifestation of antibiotic resistance. Bioorg Chem 33:149–58.

26. Brown PJB, Chang JH, Fuqua C. 2023. Agrobacterium tumefaciens: a Transformative Agent for Fundamental Insights into Host-Microbe Interactions, Genome Biology, Chemical Signaling, and Cell Biology. J Bacteriol 205:e0000523.

27. Barton IS, Fuqua C, Platt TG. 2018. Ecological and evolutionary dynamics of a model facultative pathogen: Agrobacterium and crown gall disease of plants. Environ Microbiol 20:16–29.

28. Hynes MF, Simon R, Puhler A. 1985. The development of plasmid-free strains of *Agrobacterium tumefaciens* by using incompatibility with a *Rhizobium meliloti* plasmid to eliminate pAtC58. Plasmid 13:99–105.

29. Griffitts JS, Long SR. 2008. A symbiotic mutant of *Sinorhizobium meliloti* reveals a novel genetic pathway involving succinoglycan biosynthetic functions. Mol Microbiol 67:1292–306.

30. Schnabel EL, Jones AL. 1999. Distribution of tetracycline resistance genes and transposons among phylloplane bacteria in Michigan apple orchards. Appl Environ Microbiol 65:4898–907.

31. Dai R, He J, Zha X, Wang Y, Zhang X, Gao H, Yang X, Li J, Xin Y, Wang Y, Li S, Jin J, Zhang Q, Bai J, Peng Y, Wu H, Zhang Q, Wei B, Xu J, Li W. 2021. A novel mechanism of streptomycin resistance in *Yersinia pestis*: Mutation in the *rpsL* gene. PLoS Negl Trop Dis 15:e0009324.

32. Nair J, Rouse DA, Bai GH, Morris SL. 1993. The *rpsL* gene and streptomycin resistance in single and multiple drug-resistant strains of *Mycobacterium tuberculosis*. Mol Microbiol 10:521–7.

33. Deatherage DE, Barrick JE. 2014. Identification of mutations in laboratory-evolved microbes from next-generation sequencing data using breseq. Methods Mol Biol 1151:165–88.

34. Calvopina-Chavez DG, Howarth RE, Memmott AK, Pech Gonzalez OH, Hafen CB, Jensen KT, Benedict AB, Altman JD, Burnside BS, Childs JS, Dallon SW, DeMarco AC, Flindt KC, Grover SA, Heninger E, Iverson CS, Johnson AK, Lopez JB, Meinzer MA, Moulder BA, Moulton RI, Russell HS, Scott TM, Shiobara Y, Taylor MD, Tippets KE, Vainerere KM, Von Wallwitz IC, Wagley M, Wiley MS, Young NJ, Griffitts JS. 2023. A large-scale genetic screen identifies genes essential for motility in *Agrobacterium fabrum*. PLoS One 18:e0279936.

